# Shark and ray genome size estimation: methodological optimization for inclusive and controllable biodiversity genomics

**DOI:** 10.1101/2023.02.23.529029

**Authors:** Mitsutaka Kadota, Kaori Tatsumi, Kazuaki Yamaguchi, Yoshinobu Uno, Shigehiro Kuraku

## Abstract

Estimate of nuclear DNA content serves as an independent tool for validating the completeness of whole genome sequences and investigating the among-species variation of genome sizes, but for some species, the requirement of fresh cells makes this tool highly inaccessible. Here we focused on elasmobranch species (sharks and rays), and using flow cytometry or quantitative PCR (qPCR), estimated the nuclear DNA contents of brownbanded bamboo shark, white spotted bamboo shark, zebra shark, small-spotted catshark, sandbar shark, slendertail lanternshark, megamouth shark, red stingray, and ocellate spot skate. Our results revealed their genome sizes spanning from 3.40 pg (for ocellate spot skate) to 13.34 pg (for slendertail lanternshark), in accordance with the huge variation of genome sizes already documented for elasmobranchs. Our improved qPCR-based method enabled accurate genome size estimation without using live cells, which has been a severe limitation with elasmobranchs. These findings and our methodology are expected to contribute to better understanding of the diversity of genome sizes in elasmobranchs even including species with limited availability of fresh tissue materials. It will also help validate the completeness of already obtained or anticipated whole genome sequences.

## 1. Introduction

Genome size refers to the amount of DNA in a haploid genome, expressed either by weight or number of nucleotides. Development of DNA sequencing technologies and related methods have yielded chromosome-scale sequence information with higher completeness than before, but the total number of nucleotides in resultant genome sequences rarely match the genome size, mainly because of incomplete sequencing that results from highly heterochromatic or repetitive regions in the genome. For example, the axolotl genome sequencing, a pioneering effort on a large vertebrate genome, yielded an assembly consisting of 28 gigabases^1^ (Gb), but it still lacks a large fraction larger than a whole human genome, to complete its huge genome that is estimated to amount to 32 Gb^2^.

In the early days, genome size was determined by measuring the amount of DNA extracted from a defined number of cells or nuclei. The technique revealed variable genome size among species and differential DNA contents between somatic and germ cells^3-5^. Other methods based on reassociation kinetics^6^ also provided estimations on genome size or the degree of genome complexity in various experimental settings, i.e., repetitive and non-repetitive content of the genomic DNA^7-9^. Currently, the most commonly used methods for genome size estimation are DNA staining followed by densitometry or cytometry measurement, i.e., Feulgen densitometry or flow cytometry^10,11^. These methods, however, require fresh biological samples, which is sometimes challenging, especially when the target species are elusive in the wild.

One of the taxa with scarce genome size reports is the vertebrate class of cartilaginous fishes (Chondrichthyes) comprised of chimaeras, sharks, and rays. This taxon includes some species with enlarged genome sizes over 10 picogram (pg), making it a class with a remarkable genome size variation among vertebrates^12^, next to amphibians in which genome size enlargement is observed particularly in salamanders^13^. As posed in ‘C-value enigma’^12^, what drives genome size increase/decrease remains inexplicable, and the taxon Chondrichthyes provides an intriguing system to scrutinize it in relation to the variation of habitat environment, body size, propagule size, prey, population size, and lifespan (reviewed in ref. ^14^). Among cartilaginous fishes, genome size reports are currently available only up to 134 species (based on the information in the Animal Genome Size Database: http://www.genomesize.com) among approximately 1,300 described species, which is largely due to their inaccessibility—many of them inhabit open ocean or deep-sea and thus are not immediately subjected to laboratory experiments using fresh animal tissues.

An alternative method for genome size estimation based on quantitative PCR (qPCR) was developed^15^ and is expected to circumvent the limitation of existing methods with elusive species. The method requires prior information on the genomic or transcriptomic sequences of single-copy genes to design specific oligonucleotide primers, as well as genomic DNA as a template, for PCR. Genome size is then estimated by calculating the DNA copy number contained within a defined amount of DNA. Although the qPCR-based method has been used to assess genome sizes of several plant and animal species^16-19^, its utility has been questioned. The qPCR-based method can be more labor-intensive than established methods. Moreover, in some cases, qPCR-based estimates deviated from the values obtained by other established methods such as Feulgen densitometry or flow cytometry^20,21^. Thus, establishing a streamlined protocol will allow a wide application of the qPCR-based method to provide accurate genome size estimates even in species with low sample availability.

In this study, we selected several elasmobranch species and estimated their genome sizes. We primarily adopted flow cytometry, and in parallel, we optimized the previously introduced qPCR-based method to achieve higher accuracy. This yielded genome size estimation of the species including those with low availability of fresh tissue samples, and for the brownbanded bamboo shark, the estimates were cross-validated between the two methods. Our technical optimization has been summarized in a protocol that laboratory personnel can refer at the bench for reliable genome size estimation. Based on the data we obtained, we discuss the genome size variation among elasmobranchs.

## 2. Materials and Methods

### 2.1. Animals

Red blood cells, tissue or embryo of sandbar shark *Carcharhinus plumbeus*, white-spotted bamboo shark *Chiloscyllium plagiosum*, brownbanded bamboo shark *Chiloscyllium punctatum*, slendertail lanternshark *Etmopterus molleri*, red stingray *Hemitrygon akajei*, megamouth shark *Megachasma pelagios*, ocellate spot skate *Okamejei kenojei*, small-spotted catshark *Scyliorhinus canicula*, zebra shark (or leopard shark) *Stegostoma tigrinum* (formerly *Stegostoma fasciatum*), and mouse *Mus musculus* were used for the analysis. Animal sampling was conducted in accordance with the institutional guideline Regulations for the Animal Experiments specified by the Institutional Animal Care and Use Committee of RIKEN Kobe Branch and the official guideline for animal experiments at Okinawa Churaumi Aquarium. Phylogenetic relationship of the elasmobranch species and information on the animal individuals used for the study are described in Figure 1 and Supplementary Table 1, respectively.

**Table 1.** Estimated genome sizes for species selected for this study.

| Species | Common name | Estimated C-value (pg) |  |  |
| --- | --- | --- | --- | --- |
|  |  | This study |  | Previous study |
|  |  | Flow cytometry | Quantitative PCR |  |
| *Carcharhinus plumbeus* | sandbar shark | 3.43 | - | 2.98<sup>36</sup> |
| *Chiloscyllium plagiosum* | white-spotted bamboo shark | 4.96 | - | - |
| *Chiloscyllium punctatum* | brownbanded bamboo shark | 4.83 | 5.05 | 4.56<sup>10</sup>, 4.83<sup>32</sup> |
| *Etmopterus molleri* | slendertail lanternshark | - | 13.34 | - |
| *Hemitrygon akajei* | red stingray | - | 3.82 | 4.02<sup>37</sup>, 4.15<sup>34</sup>,<br>4.32<sup>35</sup> |
| *Megachasma pelagios* | megamouth shark | - | 5.81 | - |
| *Okamejei kenojei* | ocellate spot skate | 3.40 | - | 3.09<sup>37</sup> |
| *Scyliorhinus canicula* | small-spotted catshark | 5.46 | - | 5.65<sup>33</sup> |
| *Stegostoma tigrinum* | zebra shark | 3.79 | - | - |
| *Mus musculus* | house mouse | - | 3.18 | 3.25<sup>24</sup> |

**Figure 1.**
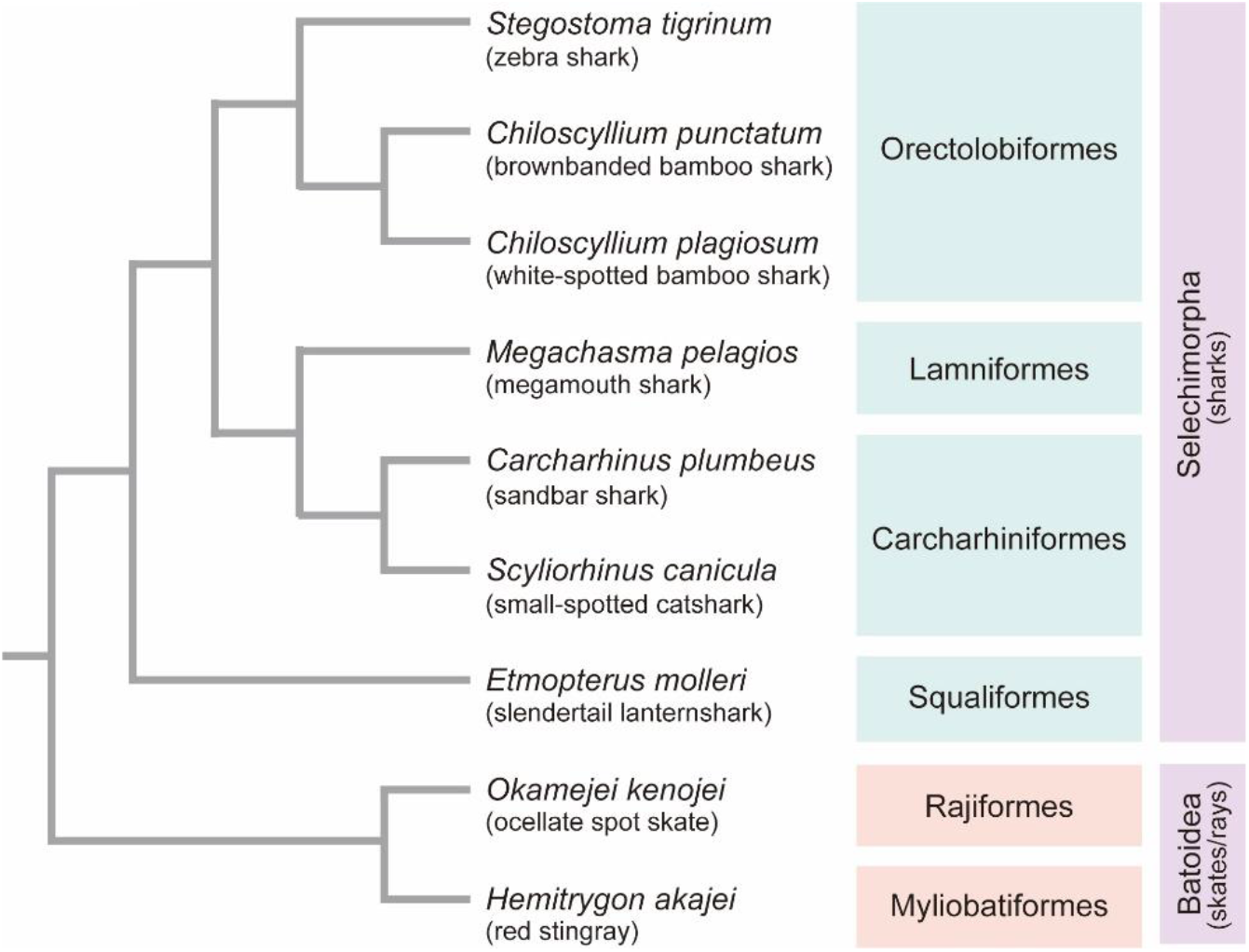
Phylogenetic relationship of the elasmobranch species used in this study. Details of the animals used in the study (e.g., source, developmental stage, and cell type) are included in Supplementary Table 1. The branch length in the tree is not scaled to relative divergence times.

### 2.2. Cell preparation for flow cytometry

Elasmobranch blood was collected using a syringe containing heparin sodium salt solution (MOCHIDA, cat: 5,000 units/5mL; use 10 μL for 2-3 ml of blood) or EDTA (0.5M EDTA, Gibco, cat: 15575-020; use 10 μL for 1-2 ml of blood). Elasmobranch cells from the tissue and embryo were prepared by mincing and dissociation with trypsin (0.5% solution in shark PBS) at 25-28°C for 15-30 min with rotation. Mouse embryonic fibroblast cells prepared from E14.5 embryos of C57BL/6 strain mice, or mouse lung fibroblast cells prepared from an adult male C57BL/6 strain mouse were cultured in DMEM (Nacalai tesque), supplemented with 10% FBS (Gibco) and 1× Antibiotic-Antimycotic solution (Gibco). Human GM12878 lymphoblastoid cell line (Coriell Institute) which has a relatively normal karyotype was cultured in RPMI (Gibco) supplemented with 10% FBS, 1× L-glutamine (Gibco), and 1× Antibiotic-Antimycotic solution. Two million elasmobranch cells or one million reference cells (mouse fibroblast cells or GM12878 cells) were permeabilized in PBS (shark PBS or PBS) containing 0.1 % Triton-X100 and stained with propidium iodide (PI/RNase Staining Buffer, BD Pharmingen).

### 2.3. Flow cytometry

Estimation of the nuclear DNA content by flow cytometry was performed as follows^22,23^. The PI intensity of the elasmobranch cells and reference cells were measured separately in a flow cytometer (Ploidy Analyser; Sysmex). Then, a single analysis setting applicable for both sample types was determined by adjusting the gain and the threshold in the CyView analysis software (Sysmex). Subsequently, the elasmobranch cells and reference cells were mixed at equal ratio and analyzed again. Genome size of the elasmobranch species was estimated relatively from the calibration curve between the 2c (G0/G1 phase) and the 4c (G2/M phase) peaks of the reference cells with known genome size information, i.e., C-value = 3.25 pg^24^ for mouse or C-value = 3.37 pg for human GM12878 cells determined by flow cytometry using mouse MEF cells as reference. The estimation is also based on the assumption that the elasmobranch species analyzed are diploid organisms. Step-by-step procedures of cell preparation, flow cytometer operation, and genome size calculation are described in the Supplementary Protocol.

### 2.4. DNA extraction for qPCR

Frozen samples (red blood cells or powderized tissue) were used for the genomic DNA (gDNA) extraction with NucleoBond AXG 100 or AXG 500 with NucleoBond Buffer Set IV (Clontech) or DNeasy Blood and Tissue kit (Qiagen). Genomic DNA was analyzed by 4200 TapeStation (Agilent Technologies) with the Genomic DNA ScreenTape analysis kit to confirm high molecular weight (HMW) DNA extraction. 1-4 μg of genomic DNA, adjusted to the total volume to 150 μL with TE buffer, was sheared to 10 kb using g-TUBE (Covaris) in an MX-300 high-speed centrifuge (TOMY) equipped with an AR015-24 rotor (TOMY) at 6,100 rpm for 1 min, at 25°C. After the shearing, size distribution of the DNA was confirmed to be around 10 kb by 4200 TapeStation with the Genomic DNA ScreenTape analysis kit. The sheared gDNA were stored at 4°C until use.

### 2.5. Selection of reference genes and primer design

Target genes for the analysis were selected from the previously defined set of Core Vertebrate Genes^25^ (CVG), consisting of 233 genes that are shared as one-to-one single-copy orthologs in the genomes of 29 selected vertebrate species. We selected the components of CVG that are also included in Core Eukaryotic Genes^26^ (CEG) and BUSCO^27^ (Benchmarking Universal Single-Copy Orthologs) vertebrate gene set identified by Hara *et al*.^25^ (see Supplementary Table S3 for detailed information of the selected genes). Orthologs of those CVGs of the target species were identified by performing BLAST (TBLASTN) search in transcriptome assemblies of the target species in the Squalomix elasmobranch sequence archive (https://github.com/Squalomix/sequences; ref. ^28^), using amino acid sequences of human CVG orthologs as the query. Also, the deduced amino acid sequences of the longest open reading frame of the identified transcripts were used for the BLAST (BLASTP) search against the human protein database. Transcripts of the target species that displayed the best hits reciprocally in the BLAST searches were considered orthologous and retained for further analysis (see Supplementary Tables S3 and S4 for the CVG transcripts used for the study). Primers were designed inside the putative last exon of the transcripts inferred by aligning the target species’ transcript sequences to the last exon of the human counterpart. Primers for the first round of PCR to generate copy-number standards were designed using the Primer3 program^29^ (https://bioinfo.ut.ee/primer3/), and primers for the nested qPCR were designed using the Primer Express software (ThermoFisher Scientific). Nucleotide and deduced amino acid sequences of CVG transcripts of the target species and the oligonucleotide sequences of the PCR primers are included in Supplementary Table S3 and S4, respectively.

### 2.6. Preparation of qPCR copy number standards

The first PCR was carried out using the KAPA HiFi HotStart ReadyMix (KAPA Biosystems) with 5 ng of the sheared gDNA and 0.3 μM concentration of each primer in 20 μL reaction volume at [98°C for 3 min, 34 cycles of (98°C for 20 sec, 58°C - 66°C in 2°C increments of temperature gradient for 15 sec, 72°C for 15 sec), 72°C for 1 min]. A portion (5 μL) of the PCR products were analyzed by electrophoresis on a 2% Agarose S gel (Nippon gene) in 1× TBE buffer and stained with GelStar (Lonza). The remaining PCR product of the condition that displayed a single-band or near single-band product on the gel was double size-selected and purified using AMPure XP beads (Beckman coulter) and eluted in 30 μL of TE. The optimal annealing temperature of the PCR reaction and size-selection condition by AMPure XP beads for each gene are included in Supplementary Table S4. Purified PCR products were sequenced directly by Sanger sequencing using the same forward and reverse primers used in the initial PCR reaction to identify possible polymorphic sites. If the polymorphic sites overlapped qPCR primer sequences, the qPCR primers were re-designed. As the first PCR was performed at a high PCR cycle number and may have reached the plateau phase resulting in the PCR product to possess a complex secondary structure, the second PCR was performed with a low cycle number using a small aliquot of the first PCR product as template. The number of PCR cycles in the second PCR was determined with QuantiTect SYBR Green PCR Kit (Qiagen) using 1 μL of the first PCR product after 5-folds of dilution or H2O as the template with 0.3 μM concentration of each primer in 10 μL reaction volume in a duplicate measurement setting, at [95°C for 15 min, 30 cycles of (94°C for 15 sec, 60°C for 15 sec, 72°C for 20 sec)] with QuantStudio 7 Flex system. The baseline signal was set to the 1st cycle only. Ct (cycle threshold) that reached 1/4^th^ of the maximum signal intensity observed at the 30th cycle of PCR was determined as the number of PCR cycles for the second PCR. The second PCR was performed using the KAPA HiFi HotStart ReadyMix (KAPA Biosystems) at the optimal annealing temperature determined earlier. The second PCR product was double-size-selected and purified using AMPure XP beads and eluted in 30 μL of TE buffer. The size distribution analysis was performed again by 2100 Bioanalyzer (Agilent Technologies) to confirm that the purified second PCR product is truly a single-band DNA. The purified second PCR product, which will be used as the DNA copy number standard (DNA standard) in the qPCR reaction, were stored at 4°C until use.

### 2.7. qPCR analysis

On the day of qPCR, DNA standards and the sheared gDNA were diluted to 1-10 ng/μL in 1× DB (dilution buffer; 10 mM Tris-HCl pH8.0, 0.05% Tween-20) using 50× DB stock solution (500 mM Tris-HCl pH8.0, 2.5% Tween-20), and 5 μL solution was used for quantitation with QubitFlex fluorometer (ThermoFisher Scientific) in four independent measurements. DNA concentration was determined by taking the average of the four independent measurements. The molecular weight (MW) of the double-strand DNA standards was calculated by the OligoCalc program (http://biotools.nubic.northwestern.edu/OligoCalc.html). The concentration of the DNA standards in copy numbers was calculated with the following formula: DNA copy number in 1 μL DNA = ((DNA concentration (ng/μL)) / (MW (g) × 10^9^)) × (6.02214076 × 10^23^/mol). DNA standards were diluted hundred-folds in 1× DB, followed by another hundred-folds dilution in 1× DB, and further diluted in 1× DB to generate 100 μL of DNA standard at 50,000 copy DNA/μL (will be STD1). STD1 was diluted five-fold in 1 ×DB by mixing 5 μL of STD1 with 20 μL of 1× DB to generate STD2 at 10,000 copy DNA/μL. Similarly, STD3, 4, and 5 at 2,000 copy DNA/μL, 400 copy DNA /μL, and 80 copy DNA /μL were generated by serial dilution. qPCR for genome size estimation was performed with the QuantiTect SYBR Green PCR Kit (Qiagen) using 1 μL of the sheared gDNA (at 1-10 ng/μL in 1× DB), STD DNAs (STD 1∼5) in 1× DB, or H2O as templates with 0.2 μM concentration of each primer in 10 μL reaction volume in 4 technical replicate setting, at [95°C for 15 min, 40 cycles of (94°C for 15 sec, 60°C for 15 sec, 72°C for 20 sec)] with the dissociation curve analysis option in QuantStudio 7 Flex system. Quantification was performed with the QuantStudio Real-Time PCR Software v1.3. The R^2^ of the standard curve and the dissociation curve were also analyzed to confirm specific amplification and quantification of the target region. C-value was determined with the following formula: C-value = amount of gDNA in the qPCR reaction (ng) / DNA copy number of the gDNA. Genome size was estimated by averaging the C-values from multiple (at least three) target gene regions. Step-by-step procedures for gDNA preparation, DNA standard preparation, qPCR, and genome size calculation are described in the Supplementary Protocol.

### 2.8. Sequence-based genome-size estimation with k-mer frequency

In parallel with the genome size estimate using the flow cytometry and qPCR, those of the zebra shark and thorny skate were estimated *in silico* based on k-mer frequency distribution with a conventional procedure. For the zebra shark, short-read genomic data produced previously by us^30^ (see Yamaguchi et al., 2022) were used. For the thorny skate, short-read genomic data produced by the Vertebrate Genomes Project^31^ was downloaded from VGP GenomeArk (s3://genomeark/species/Amblyraja_radiata/sAmbRad1/genomic_data/illumina/). Sequencing reads were trimmed with trim_galore v0.6.4 to remove low quality bases and the Illumina adaptor sequences. The processed reads (both read 1 and read 2; 351 and 751 M read pairs for the zebra shark and the thorny skate, respectively) were analyzed by jellyfish v2.3.0 program with the k value set at 21. The resultant .histo file was uploaded onto the GenomeScape webserver (http://qb.cshl.edu/genomescope/) to obtain estimates of the genome size.

## 3. Results

### 3.1. Genome size estimation with flow cytometry

Genome size estimation of nine elasmobranch species that belong to the orders of Orectolobiformes, Lamniformes, Carcharhiniformes, Squaliformes, Rajiformes, and Myliobatiformes, namely seven shark and two skate/ray species (Figure 1), was performed by flow cytometry or qPCR. The flow cytometry method utilizes propidium iodide (PI) that binds to the double-strand DNA to estimate genome sizes relative to the reference mouse or human cells with known genome size information. The reference cells at G0/G1 phase (2c) and G2/M phase (4c) were given a pre-defined nuclear genome content of 6.5 pg and 13 pg for mouse cells and 6.74 pg and 13.48 pg for human GM12878 cells to draw a calibration line. Then, the genome size of the target elasmobranch species was estimated based on the calibration line, assuming that they are diploid organisms (Figure 2). See Supplementary Table 2 for detailed information on the PI-signal values and the reference cells used in each measurement.

**Figure 2.**
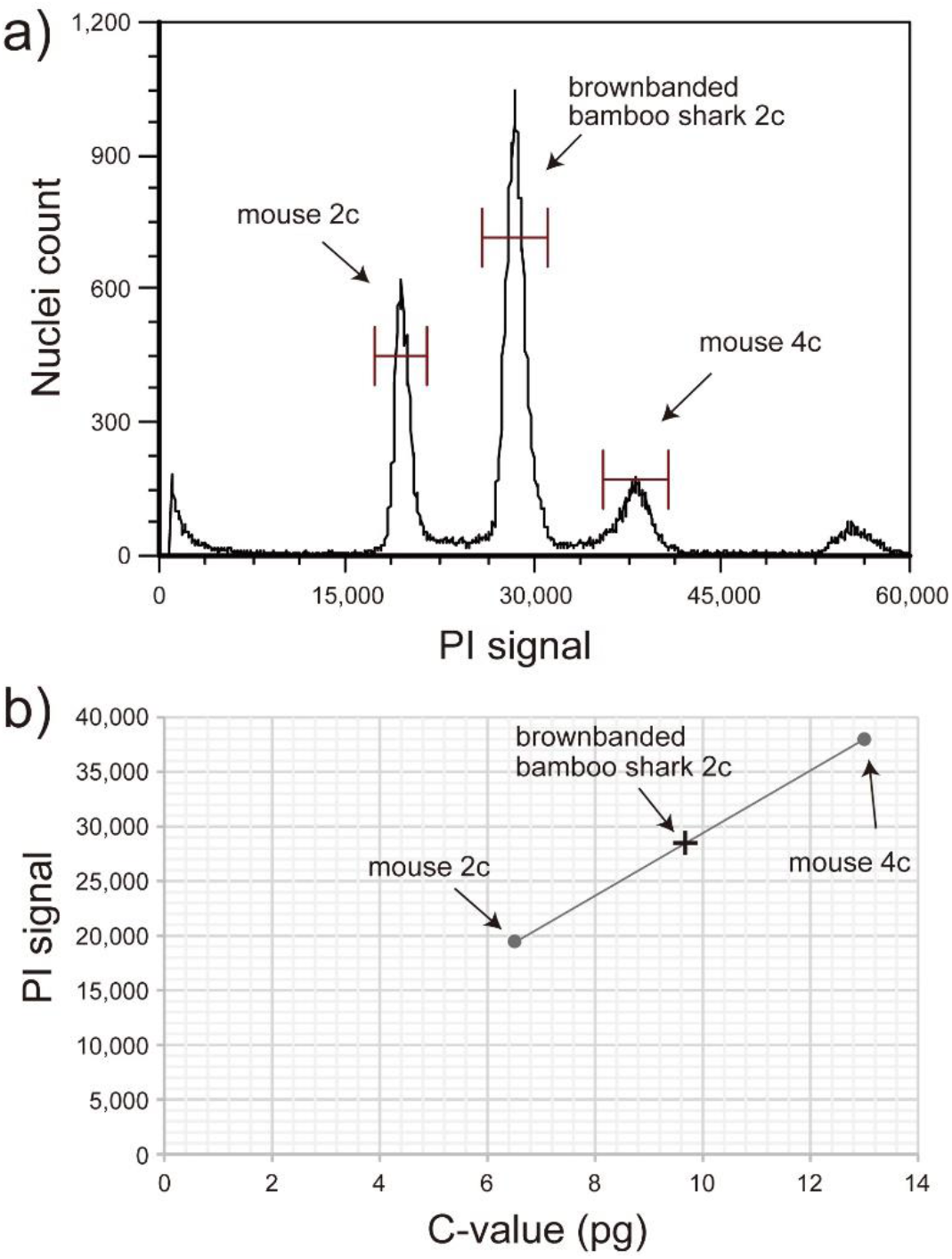
Genome size estimation by flow cytometry. a) Flow cytometry analysis of the bamboo shark *Chiloscyllium punctatum* with Ploidy Analyser (Sysmex). Red blood cells of the bamboo shark and the reference mouse lung fibroblast cells were analyzed together. PI signal of the cells corresponding to mouse 2c (cells at G0/G1 phase), mouse 4c (cells at G2/M phase), and bamboo shark 2c (cells at G0/G1 phase) were obtained. b) Genome size estimation of the bamboo shark based on the standard line of the mouse cells. C-value of the reference mouse cells was set to 3.25 pg^24^. The PI-signal values and the estimated genome sizes for the species analyzed by flow cytometry are included in Supplementary Table 2.

### 3.2. Genome size estimation with qPCR

The qPCR-based method requires the target species’ nucleotide sequence information and materials, i.e., transcriptome sequences for primer design and the genomic DNA for PCR amplification. We followed the method reported by Wilhelm *et al*.^15^ while improving the existing protocol to achieve more accurate estimation (Figure 3). The major modifications in the protocol were 1) selection of multiple (at least three) genes from the established single-copy orthologue gene set CVG, 2) use of the sheared gDNA instead of HMW gDNA for DNA quantification and PCR, 3) careful evaluation of the DNA standards; sequence analysis by Sanger sequencing and size distribution analysis by 2100 Bioanalyzer, 4) Use of a dilution buffer that contains a surfactant in the qPCR reaction. We first determined the PCR condition that gives a single-band product in the PCR reaction. The nucleotide sequence of the PCR product was confirmed by Sanger sequencing to avoid overlap of the qPCR primers with possible polymorphic sites inside the target region of the selected CVGs. The second PCR was performed using a limited number of PCR cycles, not to let the PCR amplification hit the plateau. Finally, qPCR was performed using the sheared gDNA of the target species and the DNA standards of 80∼50,000 copy/μL diluted in dilution buffer.

**Figure 3.**
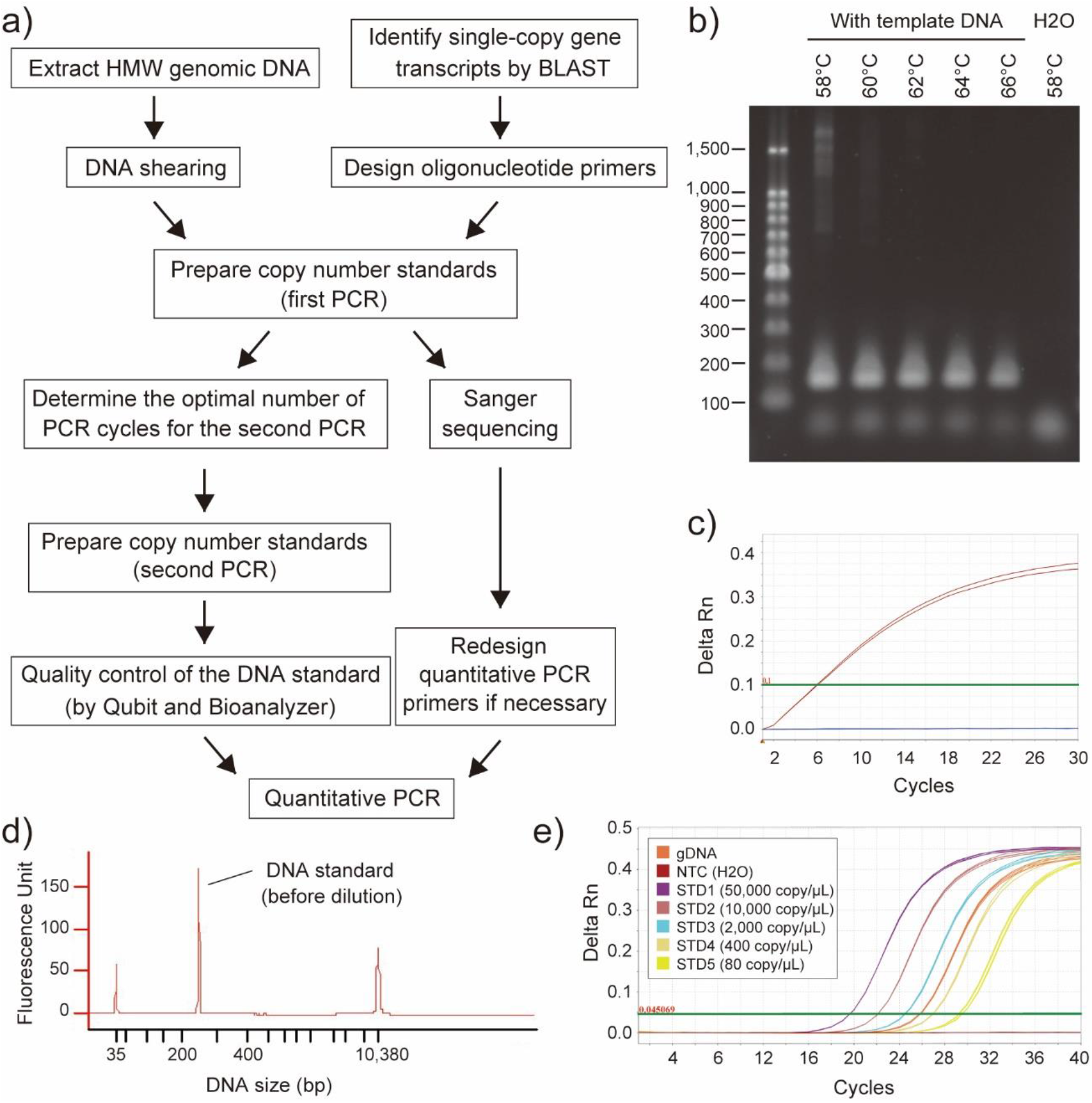
Genome size estimation by quantitative PCR. a) Workflow of the genome size estimation by quantitative PCR. b) Agarose gel electrophoresis of the first PCR product at various annealing temperature conditions. Shown is the result for the brownbanded bamboo shark MPI gene (CpMPI). c) qPCR-based determination of the PCR cycles for the second PCR. Amplification curve for CpMPI (red), negative control (H2O; blue lines), and the threshold (green line) are shown. d) Size distribution analysis of the DNA standard by Agilent Bioanalyzer with the High Sensitivity DNA Kit. Shown is the result for CpMPI. e) Amplification of CpMPI for the sheared gDNA, copy number standards (STD1-5), and negative control (NTC). Rn, normalized reporter.

The reliability of our analysis methods was assessed by comparing our estimates to the genome size information documented previously (see Table 1). The C-values of the brownbanded bamboo shark, estimated to be 4.83 pg by flow cytometry using mouse fibroblast cells as the reference (Figure 2) or 5.05 pg by qPCR, were within 5 % of difference compared to the previous documentation (4.83 pg; ref. ^32^). The C-value of the house mouse was estimated to be 3.18 pg by qPCR and was within 3 % of difference compared to the previous documentation (3.25 pg; ref. ^24^). We also confirmed that the estimated C-values of the sandbar shark (3.43 pg), red stingray (3.82 pg), ocellate spot skate (3.40 pg), and small-spotted catshark (5.46 pg) are comparable to previous reports^33-37^. For the rest of the study species selected, genome size was obtained for the first time in this study, namely for the white spotted bamboo shark: C-value =4.96 pg, slendertail lanternshark: C-value =13.34 pg, megamouth shark: C-value =5.81 pg, and zebra shark: C-value =3.79 pg. Our results of genome size estimation are tabulated under the Squalomix consortium gateway page (https://github.com/Squalomix/c-value), which will be kept up-to-date.

### 3.3. Genome sizes estimated from whole genome sequences

We also estimated genome sizes using DNA sequences with the k-mer-based method^38^ as supplementary references. For the thorny skate *Amblyraja radiata* whose nuclear DNA content was previously estimated to be 4.25 Gb^39^, the total sequence length of the publicly available genome assembly^31^ (sAmbRad1.1.pri; GCF_010909765.2) amounts to only 2.56 Gbp. Our k-mer-based estimate of its genome size using raw sequence reads resulted in an even smaller value of approximately 1.7 Gbp. Similarly, for the zebra shark *Stegostoma tigrinum* whose nuclear DNA content was estimated to be 3.70 Gb (C-value =3.79 pg), the total sequence length of the genome assembly released previously by us^30^ (sSteFas1.1; GCA_022316705.1) amounts to only 2.771 Gbp. Our k-mer-based estimate of its genome size using raw sequence reads resulted in approximately 2.805 Gbp.

## 4. Discussion

### 4.1. Importance of accurate genome size estimates—towards ‘T2T’

Recent advances in DNA sequencing technologies have enabled genome sequencing in unprecedented scale and accuracy. For setting reasonable goals in the genome sequencing project that can affect overall expenses, it is instrumental to know the karyotype and the total lengths of nucleotide sequences (namely the genome size) of the target species. Upon complete sequencing of the whole genome, the total genome sequence length should be equal to the that measured with nuclear DNA content, but in reality, the severe gap of the lengths appears especially when early technologies such as short-read sequencing is employed (see Introduction). This is the case with all cartilaginous fish species for which whole genome sequencing was conducted (Figure 4a; reviewed in ref. ^14^). In the small-spotted catshark genome assembly, more than 1 Gb, in comparison to its genome size, is missing. Also, a huge fraction (1.7 Gb, 40%) is missing in the genome assembly of the thorny skate (Figure 4). The severe gap is also often observed when genome size is estimated using k-mers, as demonstrated for the zebra shark and thorny skate in this study. As shown previously^40^, the unreliability of sequence length-based genome size estimate demands independent measurements with more reliable methods. However, the widely used methods for genome size estimation employ flow cytometry and densitometry and require fresh live materials to give an accurate estimation, which was difficult for some elusive species. To circumvent this limitation, we optimized a previously introduced qPCR-based method.

**Figure 4.**
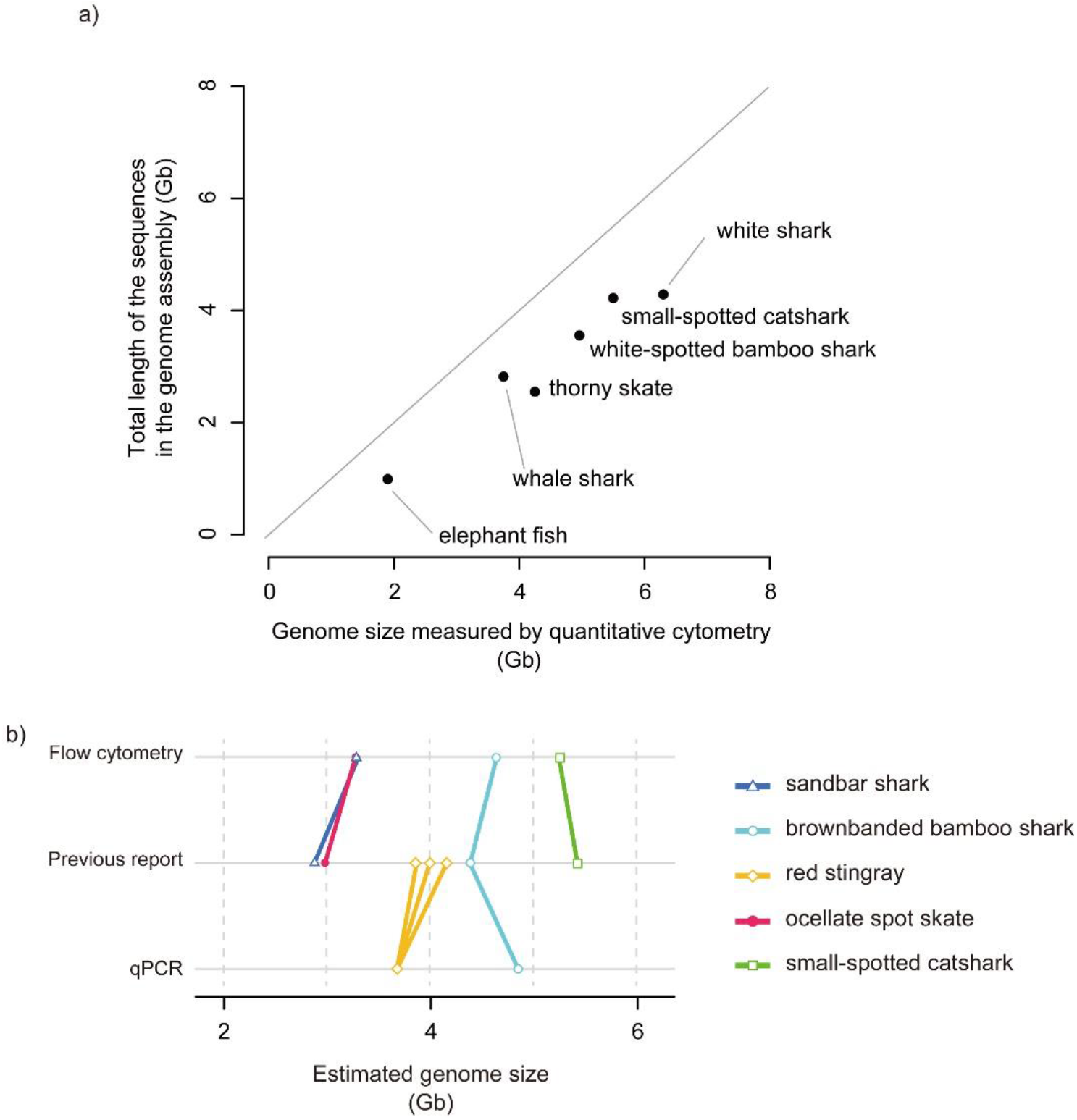
Comparisons of estimated genome sizes for cartilaginous fishes. a) Total lengths of genome assembly sequences compared with genome sizes. Included in this plot are species with unequivocal genome size estimates^32,33,39,49,50^ and whole genome assemblies^31,51-54^, namely, whale shark (NCBI Assembly ID, GCA_013626285.1), thorny skate (GCA_010909765.2), small-spotted catshark (GCA_902713615.2), white-spotted bamboo shark (GCA_004010195.1), white shark (GCA_003604245.1), elephant fish *Callorhinchus milii* (GCA_018977255.1). The genome sizes were measured with flow cytometry except for the elephant fish with the measure by Feulgen densitometry. b) Results of our genome size estimation. Estimates from previous reports are included for reference. Multiple estimates for an individual species are shown with multiple data points and lines (see Table 1).

### 4.2. Which method?—reconsideration in this post-genomic era

Our cell preparation method for flow cytometry utilized an isotonic solution (shark PBS solution)^41,42^ allowing mild permeabilization of live elasmobranch cells. Data acquisition with a flow cytometer was performed similarly to genome content analysis (e.g., ploidy level, cell cycle) using fluorescent dyes. The reference cells were cultured mouse or human cells under exponential growth conditions. Using cultured cells enabled the acquisition of PI signals for 2c (G0/G1 phase) and 4c (G2/M phase) cells to draw a regression line between these two data points. In all the measurements based on flow cytometry, we confirmed that the PI signal of the reference 4c cells is about double of the 2c cells (Supplementary Table 2), assuring that the reference data points are within the quantitative range and applicable for the estimation of relatively large elasmobranch genomes. The qPCR-based method was used for several non-shark species in past studies^16-19^. However, it received criticism about the accuracy of the estimate^20,21^. We suspected that the possible inaccuracy resulted from the lack of careful evaluation of the DNA copy number standards, careful handling in qPCR, and the use of a single target gene (instead of multiple target genes) for estimation.

### 4.3. Optimization of qPCR-based method and our protocol

In our preliminary trials, we identified several keys for improving the accuracy in qPCR-based genome size estimation and derived a protocol implementing the following modifications (see Supplementary Protocol for the entire procedure). This protocol is designated sQuantGenome (streamlined protocol for quantitative PCR-based genome size estimation) and is publicly available at the Squalomix consortium gateway (https://github.com/squalomix/c-value/). First, we used low-bind plasticwares (tubes and tips) to minimize the loss of DNA during the laboratory handling. Second, we used Qubit Fluorometer (ThermoFisher Scientific) to measure the concentration of double-stranded DNA (dsDNA) specifically. Third, we sheared HMW gDNA to approximately 10 kb-long fragments for DNA quantification (by Qubit), PCR, and qPCR. Use of unsheared HMW DNA causes inconsistent measurements in both DNA quantification and qPCR. Fourth, DNA standards were prepared with maximal caution. We designed PCR and qPCR primers within the putative last exon (that tends to be the longest among multiple exons in most eukaryotes^43,44^) of the target gene by aligning the target species’ transcript sequence to its human ortholog, based on the assumption that the exon-intron structure is conserved between elasmobranchs and mammals. The first PCR was performed at multiple annealing temperatures to identify a condition that gives a single- or near single-band product on agarose gel. We performed Sanger sequencing to confirm the identity of the nucleotide sequence of the first PCR product to the sequence from the transcriptome assembly which was obtained from the Squalomix sequence archive^28^. We also confirmed that qPCR primers (to be used for genome size estimation) do not overlap heterozygous polymorphic sites inside the DNA standard sequence. qPCR primers were re-designed if they overlapped heterozygous polymorphic sites. The second PCR was performed at a low PCR cycle number within the exponential amplification phase (confirmed by qPCR for cycle determination) to ensure that the amplified DNA is double-stranded and does not contain incomplete or chimeric PCR products. We size-selected first and second PCR products to remove DNA fragments outside the target size range. We also analyzed the DNA standards with 2100 Bioanalyzer to confirm their uniform DNA size. Fifth, we reduced the sources of variability in the qPCR reaction to improve quantification accuracy. We used a DNA dilution buffer (DB) that contains a surfactant (0.05% Tween-20) to minimize binding of the DNA to the plasticwares (tubes and tips). We prepared template DNAs (sheared gDNA and the DNA standards) in DB and quantified them with Qubit in four independent replications, using a large liquid volume (of 5 uL) containing at least 5 ng of DNA, for improved accuracy. We used an electronic 8-well pipette for consistent dispensation of the qPCR mixture into the 384-well reaction plate. We also performed the experiment with proper liquid handling techniques, e.g., pre-rinsing of the pipette tip, reverse-pipetting, etc. Sixths, our technical consideration also encompassed the *in silico* steps for selecting adequate amplification targets and primer pairs to amplify them. Choosing multiple single-copy orthologous genes (ideally no fewer than three) as amplification targets per species is a must to take a variation of estimates into consideration and to identity in inappropriate choice of genes, e.g., genes with variable copy number caused by structural variation or genes on sex chromosomes.

### 4.4. *In silico* preparation for qPCR-based genome size estimation

Recently, transcriptome sequencing with RNA-seq, as well as whole genome sequencing has led to the large-scale production of exonic sequences including protein-coding regions of various genes. This allows efficient identification of single-copy genes from the *de novo* transcriptome assembly of the target species, even when high-quality whole genome assembly is unavailable. The rigorous selection of such genes has been facilitated by the introduction of already prepared sets of single-copy orthologs, such as CEG^26^, CVG^25^, and BUSCO^27^, that were pre-selected to assess the completeness of whole genome sequence assemblies. Targeting genes from the single-copy ortholog set is a highly recommended approach for gene selection in the qPCR-based method. However, one should be aware of the possibility of lineage- or species-specific gene duplication or a gene-loss events that may cause the qPCR-based estimate to diverge from the actual genome size. In case of gene duplication, the prepared DNA standards may contain DNA of alternative sequence or alternative fragment size which should be identified with Sanger sequencing or size distribution analysis with 2100 Bioanalyzer. In case of gene loss, the candidate transcript of the target species belonging to a non-orthologous gene, may display a low similarity score or may fail to be identified as the transcript with reciprocal best hit in the BLAST analysis. In any of these cases, choosing multiple genes from the single-copy orthologous gene set and performing rigorous evaluation of the DNA standards will help prevent errors caused by the use of inappropriate genes for the analysis.

### 4.5. Genome size variation among elasmobranchs

Among the nine elasmobranch species analyzed in this study, genome size information was previously documented for five species (Table 1); sandbar shark (reported to be 2.98 pg using red blood cells with Feulgen staining^36^), brownbanded bamboo shark (reported to be 4.56 pg or 4.83 pg using red blood cells with Feulgen staining or flow cytometry^10,32^), red stingray (reported to be 4.02 pg, 4.15 pg, or 4.32 pg using red blood cells with Feulgen staining or flow cytometry^34,35,37^), ocellate spot skate (reported to be

3.09 pg using red blood cells with flow cytometry^37^), and small-spotted catshark (reported to be 5.65 pg using red blood cells with Feulgen staining^33^). Whether the estimation was based on flow cytometry or qPCR, our estimates were mostly comparable to the existing reports, suggesting its accuracy, compared with other established methods for genome size estimation. Among the species with a relatively large gap, the sandbar shark exhibited a difference of as much as 15%, between 3.43 pg (qPCR-based estimate in the present study) and 2.98 pg from a past study^36^.

We have demonstrated the utility of the qPCR-based method for genome size estimation of species whose fresh tissue material is difficult to obtain. The qPCR-based method is also recommended when suitable reference cells with known comparable genome size to the study species are not available in Feulgen densitometry or flow cytometry-based analyses, e.g., for genome size estimation of animal and plant species with extremely large genome sizes. In the case of the South American lungfish *Lepidosiren paradoxa*, the C-value was estimated to be 80.55-123.90 pg^45-47^ based on flow cytometry and Feulgen densitometry using reference cells with C-values of 3.3-6.7 pg. In the case of the canopy plant *Paris japonica* (Melanthiaceae) which has the largest genome quantified among eukaryotes to date (C-value = 152.23 pg)^48^, the genome size was estimated by flow cytometry using multiple reference cells at the C-value of 16.75 pg, 27.11 pg, and 54.08 pg. In both cases, C-value of the reference cells is much smaller than the target species, which in principle should be avoided in reference-based genome size estimation as it harbors risks of calibration error due to extrapolation. On the other hand, the qPCR method, which does not require any reference cells, is expected to provide accurate estimation for a wide variety of species with a wide range of genome sizes.

## Acknowledgment

We thank Hiroshi Kiyonari and Mayumi Watase of Laboratory for Animal Resources and Genetic Engineering RIKEN BDR for providing mouse cells, Itsuki Kiyatake and Takaomi Ito of Osaka Aquarium Kaiyukan, Kiyomi Murakumo, Rui Matsumoto, Ryo Nozu, Taketeru Tomita, and Keiichi Sato of Okinawa Churaumi Aquarium, Tomoyuki Satonaka of Shima Marineland Aquarium, Nobuyuki Higashiguchi of Suma Sea Life Park, Kazuhiro Saito, Waichiro Godo, and Tatsuya Sakamoto of Ushimado Marine Institute of Okayama University, and Takashi Asahida of Kitasato University for providing elasmobranch samples. We also thank Shinichi Nagai of Sysmex corporation for technical assistance on flow cytometry.

## Author contributions

S.K. and M.K. conceived the study. S.K., K.Y., Y.U., and K.T. collected samples. M.K. and K.T. performed experiments, and S.K. and M.K. analyzed data. S.K. and M.K. wrote the manuscript, which was approved by all authors.

## Data availability

All the details of the original data obtained in the present study have been made available in Supplementary Information.

